# Pathogenicity of Commensal Gut Biofilm In Prefrail Aging

**DOI:** 10.1101/2024.06.04.596782

**Authors:** Guillaume Le Cosquer, Melissa Pannier, Elodie Meunier, Julie Thevenin, Elise Pyhourquet, Sophie Guyonnet, Bruno Vellas, Yohan Santin, Bruno Guiard, Angelo Parini, Louis Buscail, Barbara Bournet, Damien Guillemet, Celine Deraison, Nathalie Vergnolle, Jean-Paul Motta, the IHU HealthAge INSPIRE/Open Science study group

## Abstract

Pathophysiological mechanisms of unhealthy aging, particularly the transition from robustness to frailty, remain poorly understood. Despite extensive microbiome research on taxonomy, the behavior of early prefrail gut bacteria in their natural community-host mucosal tissue context remains unexplored. Using fecal samples from the INSPIRE-T aging human cohort, we characterized gut microbiota phenotype during prefrailty stages using a polymicrobial biofilm model. Results revealed that prefrail-derived biofilms exhibited distinct taxonomic and physical alterations, enhanced dispersal, and increased epithelial virulence compared to robust counterparts. Multiparametric analyses linked biofilm characteristics to clinical traits, suggesting their potential as aging status indicators. Polyphenol-rich grape pomace extract partially reversed prefrail biofilm alterations and reduced proinflammatory prefrail biofilm responses *in vitro*. Microbiota from prefrail aged mice induced colon damage in antibiotic-treated recipients, establishing a prefrail microbiome-inflammation causality. Overall, the findings identified novel prefrail microbiome characteristics, established causal inflammatory links, and support microbiota-targeted geroprotective interventions for the prefrail populations.

## 1. INTRODUCTION

By 2030, the World Health Organization (WHO) projects that one in six individuals worldwide will be aged 60 years or older ^1^. This shift to older populations coincides with an increase in age-associated diseases and unhealthy aging. One of the first steps in the development of unhealthy aging is linked to the appearance of frailty syndrome, a clinical condition defined as decreased physiological reserve and function that heightens vulnerability and impairs the ability to withstand health stresses^2^. The absence of frailty (robust individuals) has been identified as a reliable predictor of prolonged longevity without age-associated diseases ^3^. The field of geroscience, which has recently emerged, is dedicated to understanding the underlying mechanisms of aging, finding methods to extend the healthspan – rather than lifespan – and to delay or impede frailty syndrome and age- associated diseases ^4^. The scientific community has intensely focused on describing fecal microbiome composition and changes in the aging population^5–8^. From these studies, it has become clear that chronological age is linked to alterations within the gut microbiome, a process coined as ’microb-aging’ by Bosco and Noti^6^. A current challenge is to identify additional aspects of microb-aging, beyond taxonomy, that could be related to early signs of unhealthy aging (in prefrail individuals)^9^. The involvement of microbiota behavior and virulence in such differences has not yet been investigated. Although clinical scales allow geroclinicians to categorize patients into three categories according to their frailty status (robust, prefrail, and frail individuals), biological data on the relationship between these progressive clinical changes and microb-aging dynamics are scarce. From a clinical perspective, prefrail individuals represent a potentially reversible stage before a significant decline in health. A report on community-dwelling elders indicated a notable prefrailty-to- robust transition, with a 16% conversion rate observed over an 18-to 36-month follow-up period, compared to 0% during the same period for the frailty-to-robust transition ^10^. Defining conditions or treatments that would favor this transition from prefrailty-to-robust is among the most important challenge of geroscience^10^.

The motivation of this study was thus to focus on the aging population exhibiting prefrail characteristics, in the search for interventions seeking at prefrailty-to-robust transition. For that, we capitalized on the French INSPIRE-T cohorts at Toulouse University Hospital, two clinically well-characterized human and mouse aging cohorts in which various non-invasive samples were collected, including fecal samples. These cohorts were designed to identify novel markers of aging, age-related diseases and health evolution that could be therapeutically targetable^11–13^. Considering its key importance on gut health and disease, we aimed at focusing on microbiota communities living on the surface of the mucosa, as a polymicrobial anaerobic biofilm^14^. To that end, we used a static model of polymicrobial biofilms grown on mucin-coated polystyrene microplates (MBEC device model^15–17^), providing a straightforward, mid-throughput system for parallel community analysis. Our underlying hypothesis was that these cultured biofilms, while partially preserving the fecal taxonomy, and could retain key structural and phenotypic alterations associated with both aging and donor health status. Ultimately, our goal was to establish a proof-of-concept platform for geroprotective experimental strategies aimed at addressing these abnormalities.

## 2. METHODS

### Human clinical data and samples

This study used human feces collected from the INSPIRE-T cohort in 2021 (ID NCT04224038, Table 1 and Supplementary Table 1)^12^. Donors were categorized into groups of 13 adults (7 men/6 women; 30-70 years) and 29 elderly subjects (+ 70 years). The last group was divided into 14 prefrail (6 men/8 women) and 15 healthy or robust individuals (7 men/8 women). Frailty syndrome was evaluated using the Fried frailty index (Supplementary Table 2 for detailed Fried index calculation^2^). Donors were excluded if they had received antibiotics, probiotics, or prebiotics within the past three months and if they had history of any of the following conditions: inflammatory bowel disease, irritable bowel syndrome, colorectal cancer, immunosuppression, severe liver disease. Feces were collected at the initial visit, aliquoted (≈1 g), and stored at −80°C until analysis in the INSPIRE-T biobank at a central laboratory (Toulouse Bio Ressources CRB, CHU Toulouse/IFB Purpan, Toulouse, France). Clinical characteristics, including age, gender, frailty status, and household status, were documented (Supplementary Table 1: lifestyle). For the present analysis, we used the version 1.0 of the INSPIRE-T database. We calculated the nutritional status of each individual using the Mini Nutritional Assessment (MNA; Supplementary Tables 3 and 4^18^) and calculated the Charlson Comorbidity Index based on their past medical history^19^(Supplementary Tables 5, S6, and S7). Full list of members of the IHU HealthAge INSPIRE/Open Science study group is included in Supplementary Material.

**Table 1.** Human cohort characteristics. Data are represented as mean (± standard deviation) for continuous variables and n (%) for categorical variables. MNA: Mini nutritional assessment. Statistical significance was determined by ANOVA (Fisher test) followed by Tukey’s for multiple comparisons for continuous variables and Chi squared test for categorical variables, where P<0.05 was considered significant. ^1^ Prefrail vs robust (p= 0.846) and Prefrail vs Adults (p< 0.001), ^2^ Prefrail vs robust (p< 0.001) and Prefrail vs Adults (p< 0.001), ^3^ Prefrail vs robust (p= 0.836) and Prefrail vs Adults (p< 0.001).

|  | Adults (30-70 yrs)<br>(n=13) | >70yrs robusts<br>(n=15) | >70yrs prefrails<br>(n=14) | p |
| --- | --- | --- | --- | --- |
| **Gender** |  |  |  | 0.846 |
| Male | 7 (53.8) | 6 (42.9) | 7 (46.7) |  |
| Female | 6 (46.2) | 8 (57.1) | 8 (53.3) |  |
| **Mean age (years)** | 47.8 (±11.6) | 77.8 (±5.21)<sup>1</sup> | 76.2 (±5.07) | **< 0.001** |
| **Household status** |  |  |  | 0.786 |
| Solo | 9 (69.2) | 10 (71.4) | 9 (60) |  |
| With partner | 4 (30.8) | 4 (28.6) | 6 (40) |  |
| **Mean Fried Frailty Index** | 0 (±0) | 0 (±0)<sup>2</sup> | 1.14 (±0.363) | **< 0.001** |
| **Mean Charlson Comorbidity Index** | 0.923 (±1.19) | 5.07 (±1.59)<sup>3</sup> | 4.73 (±1.87) | **< 0.001** |
| **Mean MNA score** | 27.7 (±1.58) | 28.1 (±1.27) | 27.5 (±2.30) | 0.713 |

### Mouse functional data and samples

This study involved colon tissue and feces collected from the INSPIRE mice cohort (CREFRE – US006 / Inserm – University of Toulouse, Paul Sabatier). The cohort comprised male and female SWISS outbred mice^13^. Animals underwent health assessments to measure frailty, using a rodent equivalent to the human Fried frailty index, the Valencia Score^13,20^ (Supplementary Table 8). Behavioral phenotyping was conducted by the Mouse Behavioral Core platform (Integrative Biology Center, CBI, Toulouse), at the CREFRE facility. In this study, we collected samples from young (6 months, n=8), aged robust (24 months, n=8), and aged prefrail animals (24 months, n=8). Distal colon was fixed in methacarn solution (6/3/1 of methanol/chloroform/acetic acid), and embedded in paraffin. Feces were collected and stored at −80°C. Both paraffin blocks and frozen feces were maintained in the INSPIRE mouse biobank at a central laboratory (Centre de Ressources Biologiques-Cancer - Institut Universitaire du Cancer Toulouse – Oncopole, CHU Toulouse, Toulouse, France).

### Culture of biofilms from human feces

Human fecal samples (10-20 mg) were resuspended in supplemented Brain Heart Infusion (BHI; Sigma-Aldrich 53286, St. Quentin Fallavier, France), enriched with hemin solution (final 5 µg/ml; Sigma-Aldrich H9039), Vitamin K1 (final 50µg/ml; Sigma-Aldrich V3501), and L-Cysteine (0.5 mg/ml; Sigma-Aldrich W326305). Human microbiota biofilms was grown in mucin-coated polystyrene pegs of MBEC Biofilm plates (Innovotech, Edmonton, AB, Canada) by incubating the lids in a solution of 10mg/ml gastric porcine mucin (Sigma-Aldrich M1778, a crude porcine mucin type III preparation, bound with sialic acid 0.5-1.5%) overnight at room temperature without agitation (based on methods first described in ^21^). After mucin-coating, the lids were place onto another 96-well plate containing BHI supplemented and dilution of fecal microbiota (OD 600nm =0.1; Varioskan® Flash, Thermo Scientific Massachusetts, MA, USA). Biofilms were then matured for 72 hours in a humidified incubator set at 37 °C, under anaerobic condition and continuous rocking (Rocker25 from Labnet set at 50 RPM) as described in^15–17^. After maturation, the lid was gently rinsed and basal BHI plate was refreshed to collect naturally dispersed bacteria from biofilms under the same conditions for an additional 24 hours. Biofilm biomass, viability, and matrix content were characterized using the methods described below. Additionally, the basal plate medium containing biofilm-dispersed bacteria were collected and was used for 16S rRNA V3-V4 gene sequencing, to estimate dispersal rates, and to perform interaction assays with human intestinal epithelial cells. For all microbiological procedures described herein, original sample handling and incubations were carried out in an anaerobic chamber (Don Whitley A20, 0.1% oxygen; 4.9% CO2; 95.0% N2, 45-65% humidity) or in hermetic jars with an anaerobic sachet (Anaeropack; Thermo Fisher).

### Staining assays on MBEC biofilm plates

Total biofilm biomass was evaluated using Gram’s safranin-O solution (Sigma-Aldrich), biofilm matrix-associated glycoproteins content using fluorescein wheat germ agglutinin (WGA, 1/100 diluted, ThermoFisher), and viability using a resazurin assay (1 mg/ml; Sigma-Aldrich)^15–17^. While WGA has specific affinity for sialic acid (SA) and N-acetylglucosamine (NAG), we used it to assess the presence of matrix-associated glycoproteins containing these sugar moieties (SA/NAG-glycoproteins). The dispersal rate was estimated at 24h by measuring the ratio of viable biofilm-dispersed bacteria to total biofilm biomass. Briefly, MBEC were transferred to 96 well plate containing stains and incubated at 37°C for 2 h (resazurin and WGA) or 20’ (safranin-O). Stains were recovered by sonication at 45 kHz (USC-500T, VWR) in a plate containing acetic acid (30%, safranin-O) or 1.5M sodium chloride (WGA). Specific absorbance (safranin-O) or fluorescence (WGA and resazurin) was measured using an EnSight spectrophotometer (Perkin Elmer, Villepinte, France). An average of at least 12 biofilm replicates were performed for each individual. A positive control was included for each experiment (*E. Coli* LF82, a high biofilm producer).

### Donor fecal microbiota preparation

Feces obtained from SWISS aged mice were divided into three groups of interest: young (n=8 mice, 4 males and 4 females, 6 months old), aged robust (n=8 mice, 5 males and 3 females, 24 months old), and aged prefrail (n=8 mice, 5 males and 3 females, 24 months old^13^). Feces were pooled in 1.5 mL phosphate buffered saline (PBS, Eurobio Scientific, Les Ulis, France), homogenized, and filtered through a sterile nylon mesh strainer. The filtered contents were centrifuged twice, and the pellet was resuspended in PBS/20% glycerol at OD_600nm_=1.5 (≈10^9^ CFU/ml). All the procedures were performed under anaerobic conditions at 37°C in an anaerobic chamber (Don Whitley A20).

### Post-antibiotics fecal microbiota inoculation

C57BL/6 mice (Janvier, Le Genest Saint-Isle, France), aged 8 weeks, were housed under specific pathogen-free conditions (CREFRE U006, CHU Purpan). Mice were divided into four groups: 6M (gavaged with pooled 6-month-old microbiota, n=9), 24M robust (gavaged with pooled 24-month-old microbiota, n=8), 24M prefrail (gavaged with pooled 24-month-old microbiota, n=7), and a control group receiving PBS vehicle without microbiota (n=5). Animals were housed in large cages (for rats) (maximum to 10 mice per cage) and acclimatized for 2 weeks. To control for coprophagy and potential cage effects, every two days cage bedding – including cotton nests – was thoroughly mixed every two days by a single experimenter using the same gloves throughout the procedure. After acclimatization phase, mice were distributed across four cages (PBS vehicle, 6M, 24M-robust and 24M-prefrail). The existing microbiota was perturbed by broad-spectrum antibiotics in drinking water at concentrations of 1 g/l (ampicillin and metronidazole, Sigma-Aldrich) and 0.5 g/l (neomycin and vancomycin, Sigma-Aldrich) for the first five days, and half-diluted for the remaining days. Given this standardized mixing and group distribution, we collected fecal pellets from five randomly selected animals during the acclimatization period and after antibiotic exposure. On day 10, the mice underwent fecal microbiota inoculation via oral gavage with 200 µL of previously described fecal slurry. Four days post-inoculation, the colon was harvested for evaluation of macroscopic damage, including colon thickness edema (in mm) and blind Wallace scoring endpoints^22^, was assessed by a blinded and skilled experimenter. Animal procedures were approved by the National Laboratory Animal Ethics Committee (APAFIS 2023060610225611, French Ministry of Research).

### 16S rRNA V3/V4 gene sequencing

Total DNA was extracted from feces or biofilms using the EZNA isolation kit stool DNA (VWR for biofilms) according to the manufacturer’s protocol. DNA extraction and specific PCR amplification of 16S bacterial rDNA V3-V4 regions were targeted by 341f-806R primers and sequenced on the Illumina MiSeq 2 × 250 bp technology at Vaiomer (Toulouse, France) or Biomnigene (Besancon, France). Demultiplexed raw read sequences were processed using FROGS bioinformatics pipeline (FROGS v. 4.1.0^23^, Galaxy Toulouse server, GenoToul). Adaptor and primer sequences were trimmed, and amplicons were merged using Vsearch plugin (v.2.17.0). Sequences were processed using swarm v2.0 for cluster filtering and denoising to identify amplicon sequence variants (ASVs), with a 0.005% abundance threshold. The ASV table was filtered using three criteria: minimum count of 4, 10% prevalence, and 10% variance. No rarefaction, scaling, or transformation was applied to the data. Taxonomic assignments were performed using the 16S SILVA 138.1 database.

### 16S Fluorescent In situ hybridization (FISH)

For fluorescence in situ hybridization (FISH), methacarn-fixed mouse colon tissues were embedded in paraffin. Slides were hybridized with a universal bacterial 16S fluorescent rRNA probe (EUB338-Cy3, 5’GCTGCCTCCCGTAGGAGT-Cy5, Eurofins Scientific, France) and DNA was counterstained with 4’,6-diamidino-2-phenylindole (DAPI, Invitrogen) and polysaccharide content (wheat germ agglutinin labeled with fluorescein, ThermoFisher)^16,17^. The biofilm damage score was blindly evaluated in at least three microscopic fields for each mouse in the groups of interest, according to the criteria described in^17^. Briefly, the score was established on five criteria: mucus layer invasion, bacterial-epithelial distance, biofilm density, bacterial translocation, and cell presence in the lumen. Each parameter was scored 0-3, with higher scores indicating greater mucosal biofilm disruption. Samples that were unsatisfactorily fixed for scoring were excluded from analysis. Images were acquired using a Leica LSM 710 confocal microscope (TRI Genotoul, Purpan), and FIJI freeware was used for the final image mounting (v.1.51, Mac Os).

### Human intestinal epithelial cells

Caco-2 (ATCC, Manassas, USA) and HT29-MTX E12 (ATCC) were grown separately in Dulbecco’s modified Eagle medium (DMEM, Eurobio Scientific, Les Ulis, France) supplemented with 10% fetal bovine serum (FBS, Sigma-Aldrich), penicillin/streptomycin (Invitrogen, Massy, France) and non-essential amino acid solution (Thermofisher, Paisley, UK) at 37°C in a humidified incubator with 5% CO2 as previously described ^24^. Coculture at 3:1 (Caco-2:HT29) was done on flat bottom 24-wells plates at an initial total density of 2.10^5^ cell per well (Corning, Thermo Scientific). Cells were screened for mycoplasma infection (MycoAlertTM, Lonza, Bâle, Switzerland) and treated before coculture if necessary (PlasmocureTM, Lonza). Cells were cocultured with dispersed bacteria from biofilms (normalized optical density; OD_600nm_=0.01) in OPTIMEM (Thermo Fisher). Bacterial adhesion to the intestinal epithelium was assessed after 4 h of coculture on a blood agar plate^16,17^.

### Relative mRNA expression by qRT-PCR

Total RNA was extracted using TRIzol reagent (Euromedex, France) after 4 h of coculture (Qiagen, Montreal, Canada). Reverse transcription of isolated RNA was performed using the Maxima First Strand Kit (Invitrogen, Thermo Scientific) according to the manufacturer’s instructions. Quantitative PCR (RT²qPCR Primer, Qiagen) was performed on a LightCycler (Roche, Basel, Switzerland) based on SYBR Green technology, using the Takyon Low Rox SYBR kit (Eurogentec, Seraing, Belgium). Data were analyzed by the 2^-ΔΔCT^ method using HPRT and GAPDH (Qiagen) as control genes and expressed as fold change compared to control cells without bacteria. PCR primers used in this study are listed in Supplementary Table 9.

### Grape pomace intervention

After maturation, MBEC Assay® was exposed to grape pomace solution diluted in microbiological BHI supplemented broth at several concentrations (0.3 to 30 mg/mL) for 24h. The medium was refreshed, and 4h later dispersed bacteria were collected to collect biofilm-dispersed bacteria for interaction assays with human intestinal epithelial cells. Staining assays were performed in parallel to estimate the effect of grape pomace on biofilm biomass and matrix-associated glycoproteins content. The industrial grape pomace used (VinOgrape™, Nexira, France) was an extract containing purified polyphenols (>90% content, quantified by spectrophotometry method) anthocyanins, catechin, epicatechin and oligomeric proanthocyanidins.

### Statistics and visualization

We performed statistical analyses and data visualization using R 4.3.0 – Posit (Rstudio IDE) v2023.0.9.1 built 494 (Posit Team 2023, Boston, MA, USA) and GraphPad Prism (v10 for MacOs, La Jolla, USA). Baseline clinical characteristics were compared between the groups with chi-squared test for categorical variables and ANOVA (Fisher test) for continuous variables with Tukey’s method to correct P-values. To explore the bacterial taxonomy variability among sample groups, alpha and beta diversity metrics were performed in R using phyloseq (v.1.46.0). Principal coordinate analysis (PCoA) and Partial least squares-discriminant analysis (PLS-DA) and principal component analysis (PCA) were performed in R using phyloseq (v. 1.46.0), mixomics (v. 6.26) and compared using PERMANOVA (pairwiseAdonis v.0.4.1) with 9999 permutations. To calculate microbiome uniqueness—a novel metric introduced in the context of age-related microbiome alterations^25^—we averaged the lowest beta diversity values for each individual across multiple distance metrics, including Bray-Curtis, Jaccard, unweighted UniFrac, and weighted UniFrac, compared to all other individuals within the same experimental group. This metric quantifies how distinct each individual’s microbiome is from its nearest neighbor within the cohort, with higher values indicating greater compositional distinctness. Rarefaction curves (Supplementary Figure 1A) and taxa bar plots were generated in R using phyloseq (v. 1.46.0) and ggnested (v. 0.1.0). Statistical differences between microbial communities were assessed in R using LEfSe (lefser v.1.12^26^), Deseq2 (v.1.42^27^), EdgeR (v.4.0.3^28^), metagenomeSeq (v.1.43.0^29^, and ANCOM^30^. Boxplots, barplots and scatter plots were generated in R using ggplot2 (v.3.4.4) and ggpubr (v.0.6.0). Heatmaps were created in R using complexheatmap (v.2.18.0). Grout’s test was used to filter statistical outliers in the qPCR dataset (GraphPad Prism). Statistical significance was set at P < 0.05. For group comparisons, we used Dunnett’s or Tukey’s method to correct P-values for ANOVA and the Benjamini-Hochberg method for differential abundant taxa analyses. Unless otherwise indicated, values are presented as the mean ± standard error of the mean.

## 3. RESULTS

### Characterization of the human cohort

Samples were collected from 42 individuals, stratified into 13 healthy individuals aged 30-70 years and 29 individuals aged ≥ 70 years. Within the aged group, 15 were robust based on the Fried frailty index^2^, while 14 were prefrail (score of 1 and 2) (Table 1 and Supplementary Table 1). All the donors had satisfactory nutritional scores (Supplementary Table 3). As expected, the Charlson comorbidity index ^19^ indicated more comorbidities in aged individuals than in younger donors (0.923±1.19 versus 4.73±1.87, P<0.001), but this index was not discriminant between prefrail and robust donors (5.07±1.59 versus 4.73±1.87, p=0.836; Supplementary Table 6). General characteristics of donors used in the study are summarized in Table 1.

### Taxonomic characterization of microb-aging on human biofilms

To explore links between microb-aging and prefrailty syndrome, we cultured the fecal microbiota of aged individuals on a mucin-coated device, in an attempt to reproduce a mucosal polymicrobial anaerobic biofilm^14,16,17^. Despite a decline in Bacteroidota (formerly Bacteroidetes) compared to fecal sample, biofilm communities remained dominated by Bacillota (formerly Firmicutes), specifically *Ruminococcaceae*, *Christensenellaceae*, *Bifidobacteriaceae*, *Akkermansiaceae*, *Lachnospiraceae*, and *Oscillospiraceae* (Figure 1A, Supplementary Figure 1). These taxa are core members typically found in human fecal microbiota (Supplementary Figure 2). The *ex vivo* biofilm model did not show statistically significant differences in α-diversity compared to 30-70 years individuals (Supplementary Figure 1C). Regarding β-diversity on weighted Unifrance distance, the groups were not statistically separated (PERMANOVA >0.05, Supplementary Figure 1D). However, we observed that biofilms from aged robust donors exhibited greater individual microbiome uniqueness compared to those from prefrail donors (Figure 1B). We ran various statistical models in metagenomics to identify differentially abundant taxa (i.e., Deseq2, metagenomeSeq, ANCOM, and edgeR, Supplementary Tables 10 and 11). We found correspondence in age-related taxa abundance between the biofilm model and other reports on human aging cohorts^5–8^ (Supplementary Figures 1E-F). Specifically, aged individuals presented an enrichment of taxa including *Selemonadaceae*, *Prevotellaceae*, and potential pathobionts such as *Enterobacteriaceae*. Concomitantly, there was a reduction in the abundance of important commensals within aged biofilms, including *Dialister sp., Akkermansia sp.* and *Bifidobacteriaceae* (Supplementary Figure 1E). Biofilms from aged robust individuals displayed a higher abundance of *Prevotellaceae* compared to prefrail individuals, as well as an increased representation of *Selenomonadaceae (Megamonas sp.)* and *Ruminococcaceae* (Supplementary Figure 1F).

**Figure 1:**
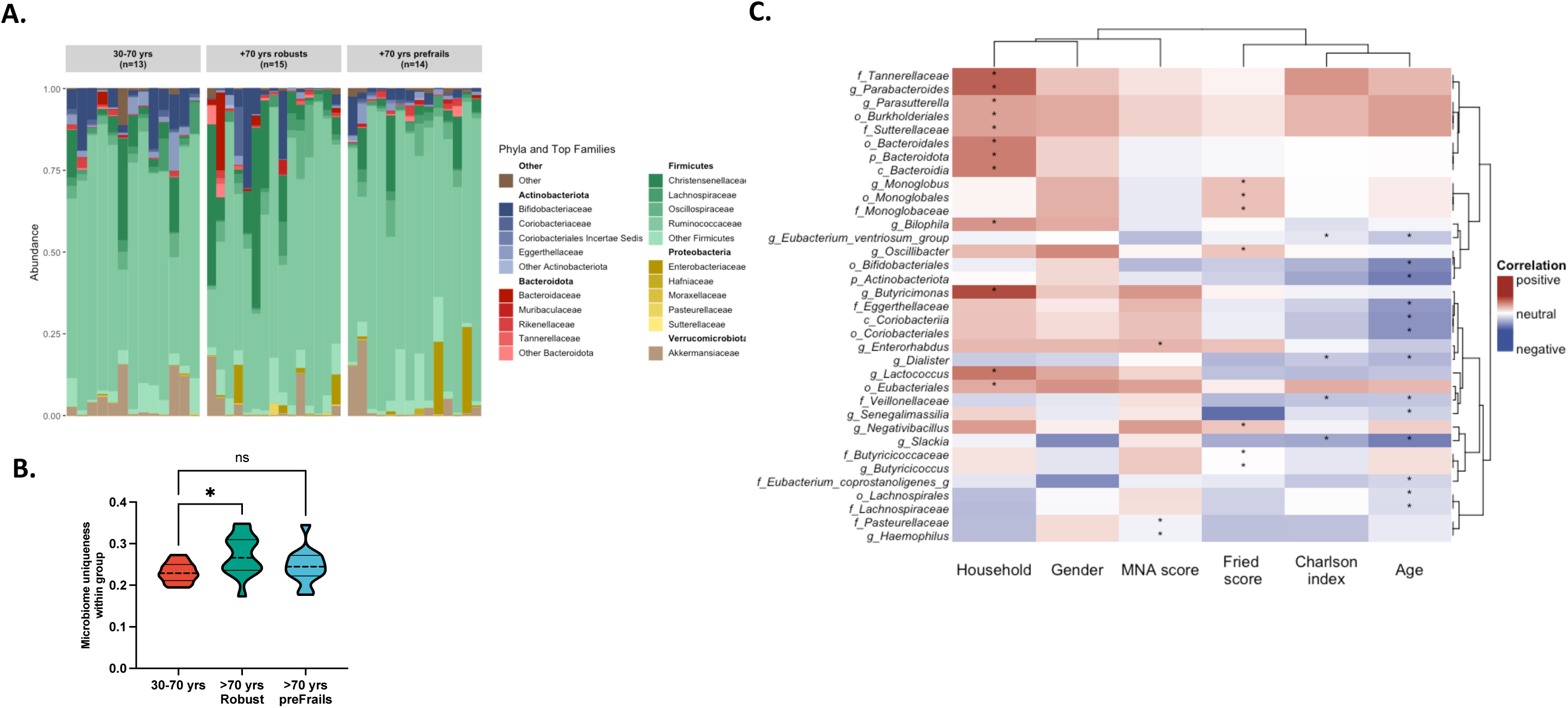
Taxonomic analyses of biofilms from human aging cohort. Fecal microbiota cultured *in vitro* on a polymicrobial anaerobic biofilm model underwent 16S rRNA V3-V4 gene sequencing. The study groups include individuals aged 30-70 years (n=13), individuals aged 70 years and above characterized as healthy aging (robust, n=15) and +70 years prefrail (n=14). **A.** The stacked bar plots illustrate the relative abundance of the 5 major Phyla and their top 5 most abundant Families. Each individual is depicted as a separate bar plot, highlighting taxonomic variations **B.** For each individual, the average of lowest values from four different beta-diversity matrices (Bray-Curtis, Jaccard, unifrac, and weighted unifrac) represented individual uniqueness. Statistical significance was determined by ANOVA followed by Tukey’s for multiple comparisons, where P<0.05 was considered significant (* for P< 0.05, *** for P<0.0010). **C.** Heatmap represents Kendall’s correlation coefficients between biofilm taxa counts clinical parameters from human donors. Clinical parameters include discrete and continuous clinical features: household (1: solo 2: with a partner), gender (1: male, 2: female), MNA score, Charlson index, age, and Fried Frailty Index. The color gradient from blue to red signifies negative to positive correlations, and * corresponds to Kendall’s P value <0.05. For enhanced clarity, the heatmap was filtered to include only significantly positive or negative correlations (P<0.05).

We then expanded the investigation between clinical characteristics of donors and taxa abundance in the biofilm model. We identified 34 statistically significant correlation pairs with biofilm taxonomic features and clinical characteristics, primarily at the Family and Genus levels (Figure 1C). First, aging alone was linked to a diminished abundance of core taxa such as *Slackia sp.*, *Eggerthellaceae*, and Bifidobacteriales. In addition, we found a positive correlation between prefrailty and the relative abundance of *Monogobales, Oscillibacter sp.* and *Negativibacillus sp* (Figure 1C). Regarding clinical history, the Charlson comorbidity index was negatively correlated with abundance of *Slackia sp*., and *Veillonellaceae (Dialister sp.)*. Abundances of *Pasteurellaceae* and *Haemophilus sp.* were inversely correlated with the MNA score, shedding light on the potential links between these specific taxa and nutritional status (Figure 1C). Gender exhibited no statistically significant correlation with the abundance of microbial taxa in cultured biofilms (Figure 1C). Finally, regarding the living conditions of donors, we uncovered a positive correlation between conjugal lifestyle and the microbial abundance of several taxa, including *Tannerellaceae*, *Parabacteroides sp.*, and *Butyricimonas sp.* (Figure 1C). Altogether, these data indicate that several microbial signatures associated with aging and prefrailty are preserved in this biofilm model.

### Physical structure of human biofilms is altered in prefrail donors

To explore microb-aging beyond taxonomy, we first characterized the physical structure of biofilms. We observed a negative correlation between donor age and total biofilm biomass (Pearson R = -0.6, P<0.05, Figure 2A). However, this feature was not significantly different between prefrail and robust biofilms (Figure 2B). Regarding the extracellular matrix of biofilms, we found a positive correlation (R=0.39, P<0.05) between age and proportion of SA/NAG-glycoproteins within this matrix (Figure 2D). Interestingly, biofilms from prefrail donors exhibited a heightened glycoproteins matrix composition compared to the other samples (Figure 2E). We then investigated the links between the clinical characteristics of human donors and biofilm physical structure (Kendall’s correlation approach, Figure 2C). In addition to confirming the previously described Fried and age relationship, we found a negative correlation between biofilm biomass and the Charlson comorbidity index (Figure 2C). Neither the donor’s sex, household, nor diet (MNAscore) demonstrated a significant association with the biofilm structure (Figure 2C). Focusing on biofilm taxonomy (Figure 2F), we identified that high biomass biofilm, a feature of 30-70 yrs biofilms, was positively correlated with a high abundance of *Eggerthellaceae*, *Oscillospira sp.*, *Enterorhabdus sp.*, and *Slackia sp.* High biofilm matrix-SA/NAG-glycoproteins, a feature of prefrail biofilms, was correlated with donors harboring low abundance of Coriobacteriales and *Eggerthellaceae* (Figure 2F). Overall, these observations suggest that the physical structure of biofilms generated from 30-70 and aged individuals differs and, more importantly, that these physical modifications may serve as discriminant features between aged individuals predisposed to prefrailty or robustness.

**Figure 2.**
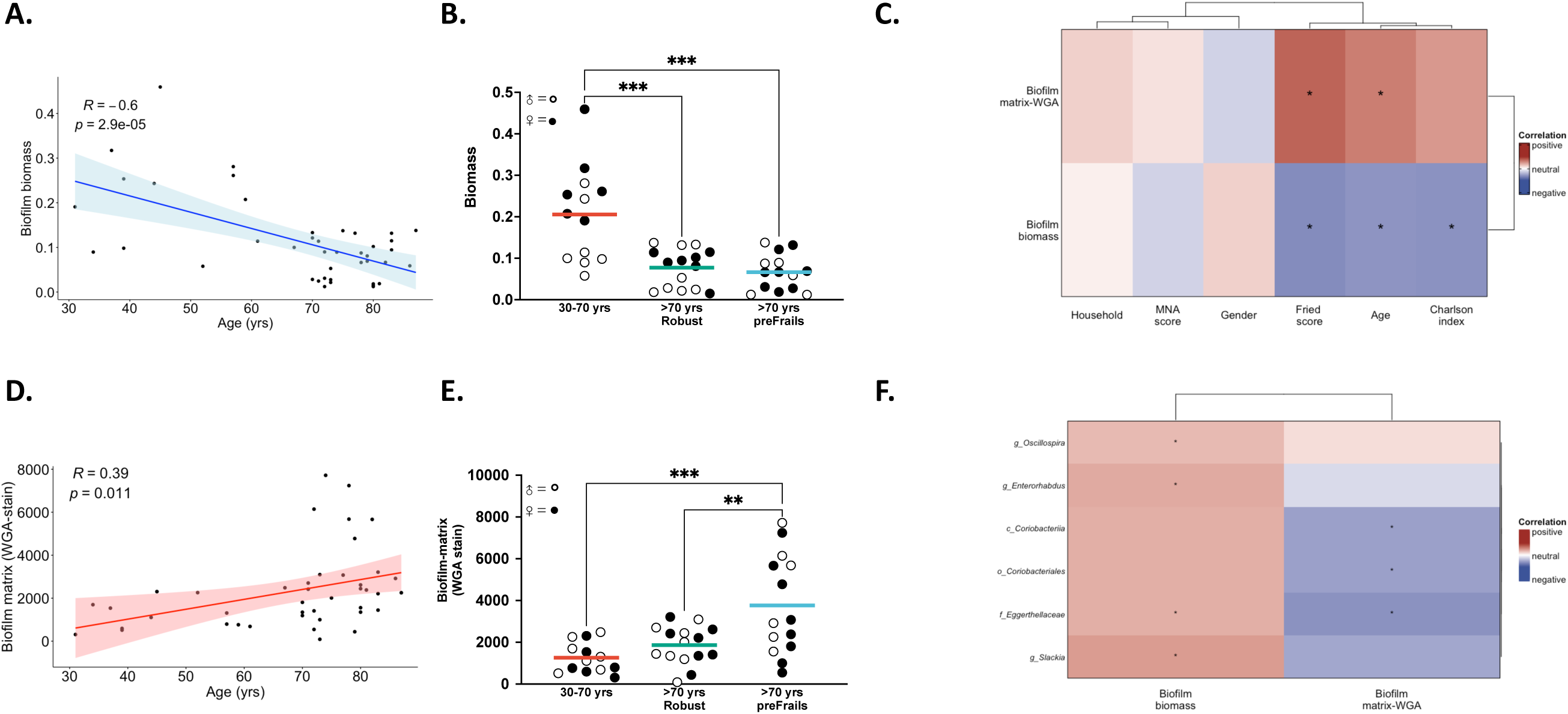
Prefrailty associated microb-aging is linked with physical alterations of human biofilms. Feces from individuals aged 30-70 years, 70+ years robust, and 70+ years prefrail, were cultured *in vitro* as a polymicrobial anaerobic biofilm. **(A-B)** The total biofilm biomass and **(D-E)** and the total sialic acid/N-acetylglucosamine glycoproteins content of extracellular biofilm matrix were measured with WGA stain. **(A-D)** Pearson correlation relationship between biofilm measures (biomass in **A.**, WGA-matrix in **D.**) and donor’s age are presented. Shaded area around the regression line represents the 95% confidence interval. **(B-E)** Scatter plots depict for each individual’s its corresponding value of biofilm biomass (in **B.**) and biofilm WGA-matrix (in **E.**) and were grouped for each three cohort of individuals. In panels B and E each dot is the average value of an individual donor biofilm (at least 12 replicates). The gender of each donor is indicated using open(male)/filled(female) circles. Statistical significance was determined by ANOVA followed by Tukey’s for multiple comparisons, where P<0.05 was considered significant. **(C-F)** Heatmaps represent Kendall (in **F.**) or Pearson (in **C.**) correlation coefficients between biofilm physical structure and clinical data (in **C.**) or biofilm taxa counts (in **F.**). Clinical parameters include discrete and continuous clinical features: household (1: solo 2: with a partner), gender (1: male, 2: female), MNA score, Charlson index, age, and Fried Frailty Index. The color gradient ranges from blue (negative) to red (positive) correlations, with statistically significant associations (P<0.05) marked with *. For enhanced clarity, the heatmap in panel **F.** was filtered to include only taxa rank with significantly positive or negative correlations (P<0.05).

### Prefrailty-associated human biofilms have an hypervirulent phenotype

In addition to the physical structure, we evaluated the pathogenic behavior of human biofilms between young and aged donors. First, we assessed the stability of biofilms, measured as the rate of bacterial dispersal over a 24-hour period. High dispersal is considered a virulent trait that increases the likelihood of exposing the subjacent host epithelium to unwanted bacterial contact ^15,16^. Donor age was not statistically correlated with this feature (P=0.058, Figure 3A). However, prefrail biofilms had a significantly higher dispersal rate than biofilms from younger groups (P<0.05, Figure 3B). We then explored the ability of biofilms to disperse bacteria that could attach to the underlying intestinal epithelium. To this end, we exposed enterocyte/mucus-producing human epithelial cell lines to a normalized concentration of the dispersed biofilm bacteria. We found that higher chronological age correlated with this behavior (P<0.05, Figure 3D). More importantly, it was dispersed bacteria from prefrail biofilms that exhibited heightened adhesiveness compared to other groups (P<0.01, Figure 3E). Intriguingly, the adhesiveness of bacteria was correlated with individuals having a high Charlson comorbidity index, suggesting a potential connection between bacterial behavior and clinical history (Figure 3C). In this cohort, neither the donor’s gender, household, nor diet (MNA score) demonstrated a statistically significant correlation’s index with biofilm pathogenic behavior (Figure 3C). Regarding taxonomy, biofilm dispersal was statistically correlated with a lower abundance of *Ruminococcaceae* and *Oscillospirales sp*. In addition, we identified several taxa whose abundance was either positively (e.g., *Adlercreutzia sp.* and Coriobacteriales) or negatively (e.g., *Akkermansiaceae*) correlated with biofilm bacterial adhesiveness (Figure 3F). In summary, our findings underscore that while chronological age plays a limited role in influencing biofilm behaviors, prefrailty status emerges as a key factor shaping the instability and virulence of gut biofilms.

**Figure 3.**
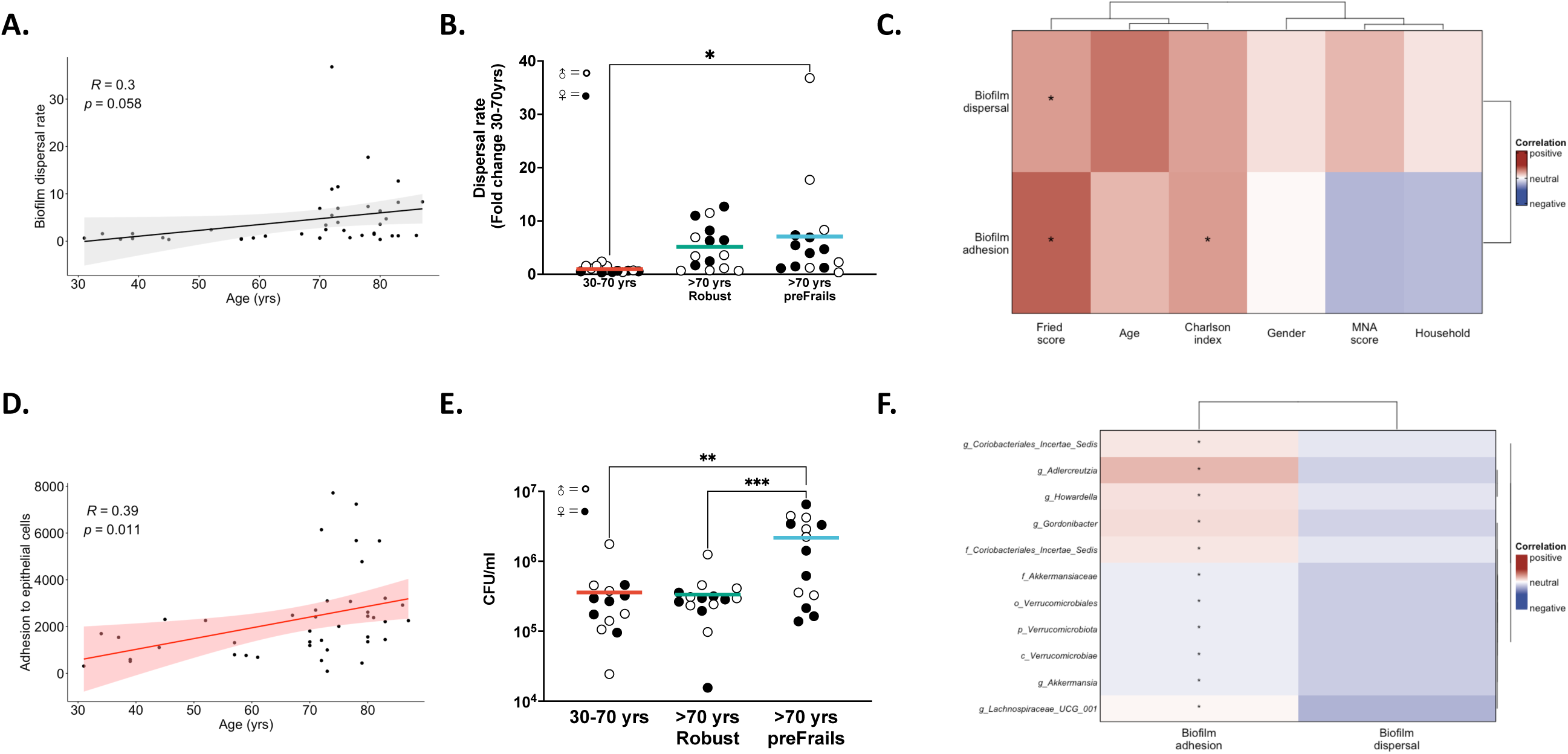
Prefrailty associated microb-aging is linked with pathogenic traits of human biofilms. Feces from individuals aged 30-70 years, 70+ years robust, and 70+ years prefrail, were cultured *in vitro* as a polymicrobial anaerobic biofilm. **(A-B)** The rate of dispersal of planktonic bacteria, biofilm-dispersed bacteria, during a 24-hour period from biofilms was measured. Raw values were normalized to the average dispersal rate of the 30-70 years group. **(D-E)** Normalized counts of biofilm-dispersed bacteria were cultured on the apical surface of the human Caco2/HT29MTX monolayer. After a 4-hour coculture, the number of bacteria adhered to the epithelium was quantified. **(A-D)** Pearson correlations between biofilm rate of dispersal (in **A.**) or adhesiveness (in **D.**) with the age of donors is presented. **(B-E)** Scatter plots depict for each individual’s its corresponding value of biofilm dispersal rate (in **B.**) or adhesiveness (in **E.**) and were grouped for each three cohort of individuals. In panels B and E each dot is the average value of an individual donor biofilm (at least 12 replicates). The gender of each donor is indicated using open(male)/filled(female) circles. Statistical significance was determined by ANOVA followed by Tukey’s for multiple comparisons, where P<0.05 was considered significant. **(C-F)** Heatmaps represent Kendall (in **F.**) or Pearson (in **C.**) correlation coefficients between biofilm pathogenic traits and clinical data (in **C.**) or biofilm taxa counts (in **F.**). Clinical parameters include discrete and continuous clinical features: household (1: solo 2: with a partner), gender (1: male, 2: female), MNA score, Charlson index, age, and Fried Frailty Index. The color gradient ranges from blue (negative) to red (positive) correlations, with statistically significant associations (P<0.05) marked with *. For enhanced clarity, the heatmap in panel **F.** was filtered to include only taxa rank with significantly positive or negative correlations (P<0.05).

#### Prefrailty-associated biofilms trigger host innate response in human intestinal epithelium

We then focused on the relative expression of 10 genes encoding proinflammatory cytokines, antimicrobials, and tight junction proteins to discern patterns indicative of the host epithelial innate response to these biofilm-dispersed bacteria. The scaled expression heatmap revealed not only altered expression patterns in biofilms from aged versus younger individuals, but also striking disparities within biofilms from prefrail versus robust individuals (Figure 4A). Proinflammatory genes, such as *IL6* and *TNF*, and the tight junction genes claudin-2 (*CLDN2*) and occludin (*OCLN*) (Figure 4B) exhibited a notable trend towards increased expression in prefrail versus robust individuals. While such trends failed to reach statistical significance when considered alone – except in the case of *TNF* and *IL6* – the PCA ordination plot demonstrated a clear shift in the transcriptome of epithelia exposed to bacteria from prefrail individuals compared to younger individuals (Figure 4C, PERMANOVA P= 0.007). As visualized in the biplot vectors, genes with the greatest contribution in the separation were those coding for proinflammatory signals (*TNF*, *IL8 and IL6* along the PC1 axis) and those related to tight junction proteins (*CLDN2* and *OCLN* in the PC2 axis). Expanding this analysis with biofilm features, we discovered that high dispersal rate – indicative of a virulent prefrail phenotype – was correlated with a higher expression of inflammatory genes (*IL6* and *TNF*, Figure 4D). Interestingly, high adhesiveness was also positively correlated with inflammatory genes (*COX2*) as well as genes encoding tight junction proteins (*CLDN2* and *OCLDN*; Figure 4C). Finally, we found that an increased quantity of SA/NAG-glycoproteins in the biofilm matrix – a feature linked to aging and prefrailty – was positively correlated with a higher expression of proinflammatory genes (*IL6*, *IL8*, and *TNF*). Overall, these findings indicate that in addition to aging host immunity, microbiota biofilms from +70 years prefrail individuals, but not in 30-70 years and +70 years robust individuals, contain commensals with proinflammatory triggering potential.

**Figure 4.**
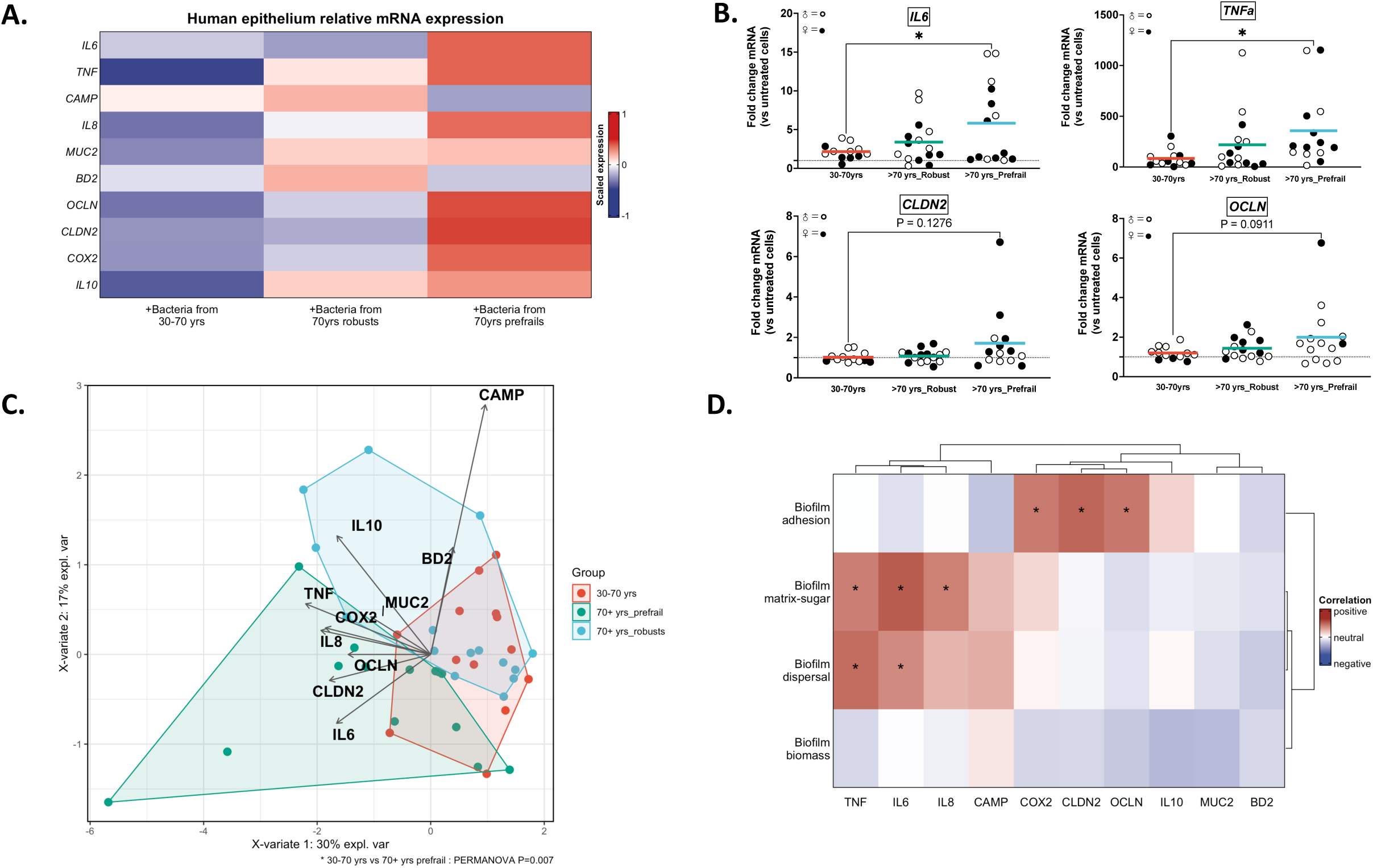
Increased host epithelial inflammatory response to prefrailty-associated human biofilms. After coculture with a normalized amount of biofilm-dispersed bacteria, epithelial cells were collected for RNA extraction. **A.** The heatmap represents scaled relative expression of a targeted list of host response genes. Specifically, these genes are involved in innate inflammatory signaling (*IL6*, *TNF*, *IL8*, *IL10*, *COX2*, and *BD2*), secretion (*BD2* and *MUC2*), and tight junction function (*OCLN* and *CLDN2*). The color gradient from blue to red correspond to scale relative mRNA expression. **B.** Scatter plot depicts relative mRNA expression of *IL6*, *TNF*, *CLDN2* and *OCLN* for each individual’s and were grouped for each three cohort of individuals. The gender of each donor is indicated using open(male)/filled(female) circles. Dotted line at 1 corresponds to basal mRNA expression in wells without bacteria. Statistical significance was determined by ANOVA followed by Tukey’s for multiple comparisons, where P<0.05 was considered significant. **C.** Gene expression dataset was ordinated using Partial least squares-discriminant analysis (PLS-DA) with its corresponding loading vectors. Pairwise PERMANOVA revealed statistical differences between the group of 30-70 years and +70 years prefrail (P=0.007). **D.** The heatmap corresponds to Pearson correlation coefficients between host epithelial gene response and biofilm features. The color gradient from blue to red signifies negative to positive correlations, and * corresponds to Pearson P value <0.05.

#### Prefrailty aging in mice is linked to altered mucosal biofilm biogeography in colon

Motivated by human biofilm model findings, we aimed to confirm whether early signs of prefrailty could influence the spatial organization of the gut microbiota in the colon. For this, we used tissues from an aging mouse cohort, equivalent to humans aged 30 to 70+ years (= 6 to 24 months old, INSPIRE mouse cohort^13^). Among the 15 selected animals, thorough frailty syndrome assessments were conducted, categorizing them into groups mirroring healthy aging (robust, n=8) and those displaying early signs of frailty (prefrail, n=7, Valencia Frailty Score, Supplementary Table 8^20^). Upon sacrifice, the colon-associated microbiota was visualized using 16S fluorescent *in situ* hybridization (FISH) staining (Figures 5 A-C). Remarkably, young mice exhibited a mucosal microbiota organized as a dense biofilm community lining a layer of sterile mucus (outlined by dashed lines in yellow, Figure 5A). In contrast, aged animals displayed a different spatial organization, featuring isolated bacteria within the mucus layer that escaped from the dense mucosal biofilm (arrows, Figures 5B and 5C). Bacterial translocation, which was almost absent in the young cohort, was observed in the aged animals (Figures 5B and 5C). Notably, aged prefrail animals exhibited more signs of tissue-associated biofilm alterations than aged robust animals, including signs of virulent behavior in gut microbiota such as invasion of the mucus sterile layer, epithelial adhesion, or translocation (biofilm damage score^17^, Figure 5D). These findings confirmed *in vivo* that prefrail status in aging is linked to altered biofilm organization and emergence of virulent phenotypes.

**Figure 5.** Prefrailty aging in mice is linked with altered gut biofilm and emergence of pathogenic behavior when transferred into recipient mice. **(A-C.)** Using 16S fluorescent *in situ* hybridization, we visualized tissue-associated microbiota in the distal colon of young (6 months, 6M, n=8, **A.**), aged (24 months, 24M) robust (n=7, **B.**), and aged prefrail (n=7, **C.**) mice. Representative images are presented with blue is DAPI staining for host nuclei, green is fluorescein-coupled wheat germ agglutinin for glycoproteins-rich content (e.g., mucus layer), and red is the 16S-Cyanine3 probe for all bacteria. Dashed lines represent the limits of mucosal tissue and the "inner mucus layer". Arrows highlight areas of interest. Scale bars are 25 µm. **D.** Biofilm-damage scores were calculated for the three groups of mice, with dots representing the mean score from at least 3 fields per animal. **(E-G.)** C57Bl/6 animals (8 weeks old) were previously exposed to 10-day antibiotics and underwent oral gavage with either vehicle (PBS, n=5), or with feces from young (6M, n=9), aged robust (24M robust, n=8), and aged prefrail (24M prefrail, n=7) mice. **E.** Four days after inoculation, feces were collected for microbial taxonomy assessment using 16S rRNA V3-V4 gene sequencing. Colons were harvested for (**F.**) parietal thickness measurement and (**G.**) macroscopic damage score. **(D, F, G)** Bar plots corresponds to mean value per group ±SEM. Statistical significance was determined by ANOVA followed by Dunnett’s for multiple comparisons, where P<0.05 was considered significant (* for P<0.05, ** for P<0.01).

#### Prefrailty-associated microbiota directly causes colon damage when inoculated into young mice

To experimentally demonstrate a causal relationship between prefrailty-associated microbiota and local gut inflammation, we orally inoculated fecal microbiota in antibiotic-treated young mice. We used as inoculate a pool of feces from young (6 months, n=8), robust (24 months, n=8), and prefrail mice (24 months, n=8). As expected, the antibiotic regimen caused taxonomic dysbiosis in gut microbiota, with severe reduction of total load, alpha diversity, and an increased abundance of *Enterobacteriaceae*, which was partially restored after 4 days of antibiotic washout (Figure 5E, Supplementary Figures 3-5). Post-inoculation, animals were monitored daily for 4 days, and no animals reached a distress endpoint (Supplementary Figure 6A). Mice inoculated with prefrail microbiota exhibited a significant difference in beta-diversity compared to those receiving aged robust or young microbiota (Supplementary Figure 4D). Furthermore, we observed a higher abundance of Proteobacteria, and a reduced abundance of *Muribaculaceae*, *Christensenellaceae*, *Parabacteroides* sp., and *Akkermansia* sp., in mice inoculated with prefrail microbiota compared to the other groups, mirroring previous findings in the litterature^31^ (Supplementary Figure 5). Regarding intestinal tissue damage after the inoculation procedure, we found that young fecal microbiota did not cause macroscopic harm in the colon (total damage score of 1.7±0.44, Figure 5F-G, Supplementary Figure 6B). When inoculated with aged prefrail microbiota, recipient animals had significantly higher colon damage such as increased incidence of erythema, mucus release in the lumen, edema, and (total score 5.4±0.98, ANOVA followed by Dunnett’s test, P = 0.002, Figure 5G). Additional histological analysis on these mice using FISH staining showed reduced bacteria-epithelium distance in prefrail mice compared to young mice (but not compared to aged-robusts), indicating a potentially compromised colonic barrier integrity in the prefrail group (Supplementary Figure 6C). These results combined with previous *in vitro* assays on intestinal epithelial cell lines, suggest that microbiota from prefrail – but not from young and robust – may directly causes intestinal damage when inoculated into recipient young mice.

#### Grape pomace partially reverses prefrailty-associated biofilm structure and virulence

Considering the virulent phenotype of the prefrail microbiota and its associated damage to the intestinal mucosa, we investigated whether this phenotype can be reversed by microbiota-targeted strategies. Grape-derived polyphenols have demonstrated prebiotic properties via their activities on gut microbes^32–35^. Indeed, the majority of edible grape polyphenols are poorly bioavailable and reach the colonic compartment where they shape microbiome profile^36^. Therefore, we sought to determine whether industrial grape pomace, which is already marketed as a food supplement in France, can affect the structure and virulence of mucosal biofilms associated with prefrail aging. We previously observed that prefrail biofilms had reduced biomass compared to young biofilms, and exposure to grape pomace significantly increased it (Figure 6A, P<0.05 at 30 mg/ml). Importantly, we found a statistically significant reduction of 79.5% in the dispersal rate in prefrail biofilms (Figure 6B, P<0.05 at 30 mg/ml). However, grape pomace had no significant effect on the increased adhesiveness of dispersed bacteria from prefrail biofilms (Figure 6C). At a concentration of 30 mg/ml, biofilm exposure to grape pomace led to a significant reduction of dispersed bacteria able to trigger the expression of *IL6* (55% reduction, P<0.05, Figure 6D), while having no statistically significant impact on levels of *IL8* and *TNF* (P>0.05, Figure 6E-F). Overall, these experiments using a biofilm model suggested that grape pomace could directly reduce the hypervirulent phenotype associated with the prefrail aging microbiota.

**Figure 6.**
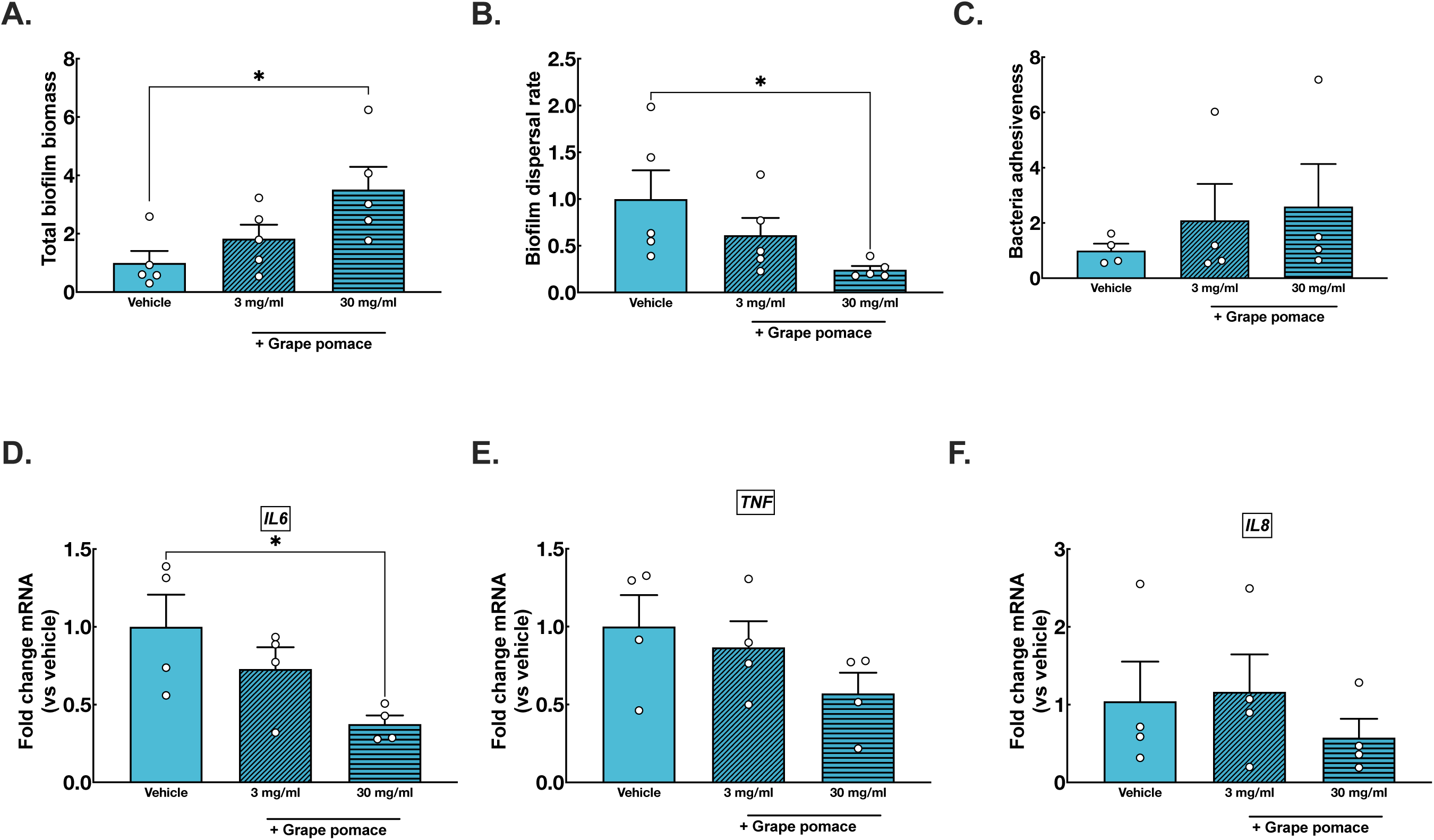
Positive impact of grape pomace on prefrailty-associated human biofilm structure and virulence. Polymicrobial anaerobic biofilms were generated from human feces of prefrails individuals aged >70yrs, n=4-5 per group) and exposed to grape pomace diluted in microbial broth at concentrations of 3 to 30 mg/ml for 24 hours. **A.** Total biomass was evaluated using safranin-O stain, and the data are presented as the fold change of mean biomass from control prefrail biofilms (broth media). **B.** The rate of dispersal for each biofilm was evaluated as a ratio between amount of biofilm-dispersed bacteria and biofilm biomass previously assessed. **C.** Normalized counts of biofilm-dispersed bacteria were cultured on the apical surface of the human Caco2/HT29MTX monolayer. After a 4-hour coculture, the number of bacteria adhered to the epithelium was quantified by plating and was represented as fold change of vehicle treated biofilms (broth media). **D-F**. Dispersed bacteria were collected from biofilms previously exposed to grape pomace. Human intestinal epithelial cell lines (Caco2 and HT29MTX, at a 3:1 ratio) were then exposed to these bacteria for 4 hours, and mRNA expression of *IL6* (**D.**), *TNF* (**E.**) and *IL8* (**F.**) genes were assessed. **(A-C)** Bar plots corresponds to average value ± SEM of 12 biofilm replicates from human donors (represented as individual dots). **(D-F)** Bar plots corresponds to average value ± SEM of relative mRNA expression of vehicle control group (n=4 human donors). Statistical significance was determined by ANOVA followed by Dunnett’s for multiple comparisons, where P<0.05 was considered significant and * for P<0.05 and *** for P<0.001.

## 4. DISCUSSION

Aging is commonly associated with changes in the gut microbiota, including shifts in taxonomy and metabolism, a phenomenon referred to as microb-aging ^6^. Despite extensive research on fecal microbiome from various populations worldwide (reviewed in^5–8^), significant knowledge gaps remain on the ecological implications of microb-aging within commensal gut communities. Another under-explored area is the causal relationship between microb-aging and low-grade inflammation of aging ^9,37^. Additionally, the role of microbiota in frailty aging and the potential of microbiota-targeted therapies in transitioning from prefrailty to robustness remain open questions. Addressing these gaps is crucial for understanding the nature of microb-aging and identifying non-chronological markers of unhealthy aging ^4,11^. Using non-invasive samples in a clinically well characterized human aging cohort, we thus aimed at generating a model for microbiota forming biofilm, which is the microbial lifestyle that resides in close proximity to the host tissue and epithelium ^14^. In this study, we aimed at detecting microbiome shifts in biofilms from aged individuals at an early stage of frailty, including but not limited to taxonomy, specifically to uncover potential therapeutic interventions that can reverse these changes and establish new targets that promote the transition from prefrailty to robustness ^10^.

From a taxonomic perspective, the cultured biofilm maintained important microb-aging features already described in the literature, such as increased microbiome uniqueness in robust but not in prefrail aged individuals (Figure 1B)^25^. By comparing biofilm taxonomy results with a recent grouping proposed by Ghosh et al.^5^ — Groups I, II, and III—we observed an interesting parallelism, where gut pathobionts, such as Proteobacteria from the *Enterobacteriaceae* family, exhibited increased abundance in aged individuals. We also noted a decline of beneficial taxa in aged biofilms, such as *Bifidobacteriaceae*. Interestingly, *Akkermansia sp.*, a genus containing important commensals involved in host/mucus homeostasis^38^, showed slightly reduced abundance in aged biofilms. Intriguingly, biofilms from aged robust individuals displayed increased taxa traditionally associated with cohorts of healthy centenarians such as *Christencellaneaceae* ^39–41^. As a limitation of these taxonomy findings relates to small sample size, further investigation within the entire INSPIRE cohort could be engaged to confirm and refine the identified taxonomic features (+1000 volunteers^12^). In addition, the longitudinal design of this cohort will enable collection from the same individuals over the next 10 years, enabling us to track the progression of prefrailty in individuals and determine whether they experience a shift to frailty, provided that microb- aging is not therapeutically managed (with diet, exercise or pre- and probiotics).

In addition to taxonomic alterations, this study revealed new microbiological features that may help distinguish between healthy and unhealthy aging in humans. Specifically, we identified a reduced biofilm-forming capacity in most fecal samples of aged individuals compared to that in younger individuals (Figure 2). This may indicate a common biological pattern in the aging microbiome that is potentially linked to the loss or gain of specific taxa, as previously discussed. More compelling are those features that distinguish between aged- matched robust and prefrail donors, such as the amount of sialic acid/N-acetylglucosamine-- rich glycoproteins associated with the biofilm matrix (Figure 2). Extracellular polysaccharides are indeed fundamental for retaining water within the biofilm matrix, generating a highly hydrated and cohesive environment that influences cell movement ^42,43^. Whether the altered glycoproteins composition of the aged prefrail biofilms used in the present study contributes to taxonomic changes, increased dispersal, and fragmented biofilm physical structure *in vitro* (Figure 3) and *in vivo* (Figure 5) remains to be experimentally elucidated. Another discriminant feature of prefrailty-associated biofilms is their capacity to release bacteria with higher adhesiveness and which triggers proinflammatory signals from the underlying epithelium (Figure 4). Although this phenomenon has not been previously documented, metagenomic studies of fecal microbiota have indicated an increased proportion of virulence gene pathways in microbiota from aged frail individuals ^44^.

It is yet to be fully understood whether local gut microbiota alterations directly contribute to adverse outcomes in aged frail populations. Complementing *in vitro* findings, an *in vivo* microbiota inoculation approach reinforces the hypothesis that prefrailty associated microbes could directly cause a sub-inflammatory phenotype in the gut. Although this aligns with other studies of aged microbiota transfer ^45^, the novel message from the present study lies in the separation between robustness and prefrailty-associated sampling. Here, we found that macroscopic colon damage induced by aged microbiota may be more pronounced with prefrail microbiota than with robust (Figure 5G). Undoubtedly, the interaction between the gut microbiome and its host is bidirectional and more complex links are expected *in vivo*. A road to be explored is the impact of a prefrailty microbiota within an aging host, where heightened inflammatory stress and immunosenescence may compromise the host’s ability to mount an effective response to local insults (the inflammaging hypothesis ^37^). In addition to the virulent behavior of prefrail biofilm, local inflammation may significantly alter microbial biotope making it less favorable to commensal gut bacteria, or even promoting the blooming of pathogenic taxa (e.g., *Enterobacteriaceae* ^46^). This could be attributed to compromised epithelial secretory functions in high age, such as defective mucus and antimicrobial peptide production^47^, or deficiencies in intestinal barrier integrity ^48^– although the latter phenomenon remains a subject of debate ^49^. Natural polyphenols, including those present in grape pomace, have been shown to improve cognitive function, extend the lifespan, and protect against age- related muscle loss in several mice models ^32–35^. Studies in mice have shown that grape pomace in the diet can improve metabolic diseases by preventing gut microbiota dysbiosis and increasing the abundance of beneficial taxa ^50–52^. In this study, we found that an industrial grape pomace preparation favored the restoration of the physical and non-pathogenic features of prefrail biofilms *in vitro*, potentially indicating the improved resilience of microb- aging communities (Figure 6). Additionally, this compound reduced the release of proinflammatory triggering bacteria from prefrail biofilms. Previous study suggested that grape pomace did not cause a significant taxonomic shift in human gut microbiota ^53^. Therefore, further research is needed to determine the precise impact of grape polyphenols on microbial virulence functions and to elucidate specific microbial genetic responses. Finally, going beyond this particular compound, the methodology presented here may support the clinical relevance of gut microbiota biofilm cultures (using non-invasive fecal samples) to assess and validate the effects of new classes of geroprotectors, such as prebiotics or probiotics, specifically targeting signs of unhealthy microb-aging.

Beyond the well-known taxonomic changes in the aging gut microbiome, this study points to novel functional and behavioral changes in human gut biofilm, that switched to a virulent phenotype in prefrail individuals but not in robusts. Importantly, we found that this phenotype was reversible by polyphenol-rich grape pomace treatment. Altogether, these results further establish the intestinal microbiota as a target to improve healthy aging. These discoveries might represent a unique opportunity for geroclinicians to develop screening tools and identify new therapeutic targets against microb-aging, to prevent frailty development in the expanding aging population.

## Author Contributions

GLC. performed most experiments, analyzed data, and provided clinical input; MP., EM., JT., EP., CD., NV. and JPM. contributed to *in vivo* and *in vitro* data acquisition and analysis; SG., and BV. provided human samples, contributed to clinical characterization, and reviewed manuscript; YS., BG., and A.P provided mice samples, contributed to clinical characterization and reviewed manuscript; LB., BB. reviewed manuscript and provided clinical input; DG. provided scientific reagents and reviewed the manuscript; JPM. performed 16S bioinformatics and statistics; GLC., NV. and JPM., wrote the manuscript; N.V and JPM. supervised the work and obtained fundings. All authors have read and agreed the final version of the manuscript.

## Data Availability

16S rRNA gene sequencing from the experimental biofilms and mouse data were uploaded to the NCBI Sequence Read Archive (SRA Bioproject PRJNA1076354). Additional biological or clinical metadata from the INSPIRE-T cohort can be obtained by contacting the corresponding author, who will submit the request to the INSPIRE biobanking and research committee for final approval. Other information from this study can be obtained, upon reasonable requests, from the corresponding author.

## Ethics

The INSPIRE cohort protocol ID NCT04224038 was approved by the French Ethical Committee located in Rennes (CPP Ouest V) in October 2019. Mice experimentation protocol has been approved by local ethic committee of Toulouse University and by the Ministry of Superior Education and Research under the agreement APAFIS2023060610225611.

## Acknowledgments

Authors thank the Department of Gastroenterology and Pancreatology at the University Hospital of Toulouse for an MD/PhD scholarship leave to GLC. Authors give credit to Dr. Felipe Sierra for its initial assessment of the research project, under the umbrella of INSPIRE Call for Geroscience Research Grant. Authors thank the INSPIRE biobanking committee for evaluating our project and approving sample collection. Authors thank the staff of the human and mice Centre de Recherche Biologique (CRB) of the Toulouse Hospital for samples preparation. Authors thank the CREFFRE core facility (UMS 006) for hosting animals used in the project and contributing to histopathology analyses (R. Balouzat and S. Milia), and the TRI-Genotoul platform for cellular imaging facilities (U1043, S. Allart). Authors are thankful to Dr. Racaud-Sultan for critically reading the manuscript. Fundings came from INSPIRE-Grant AMIGO / ANR dysBIOFILM ANR-22-CE14-0041-01 (JPM), ANR-PARCURE PRCE-CE18, 2020 (NV), and the national program "Microbiote" from INSERM (NV and JPM). The INSPIRE-T study was supported by grants from the Region Occitanie/Pyrénées-Méditerranée (Reference number: 1901175), the European Regional Development Fund (ERDF) (Project number: MP0022856), and the Inspire Chairs of Excellence funded by: Alzheimer Prevention in Occitania and Catalonia (APOC), EDENIS, KORIAN, Pfizer, Pierre-Fabre. The IHU HealthAge Open Science initiative was supported by the French National Research Agency (ANR) as part of the France 2030 program (ANR-23-IAHU-0011), and builds on the work conducted in the Data Sharing Alzheimer project.

